## Supplemental Figures and Tables for "CLM296: a highly selective inhibitor targeting ALDH1A3-driven tumor growth and metastasis in breast cancer"

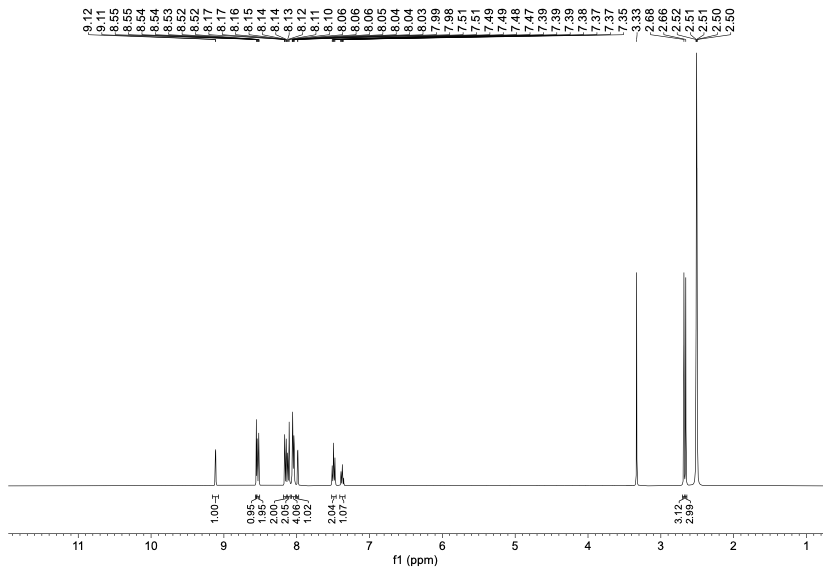

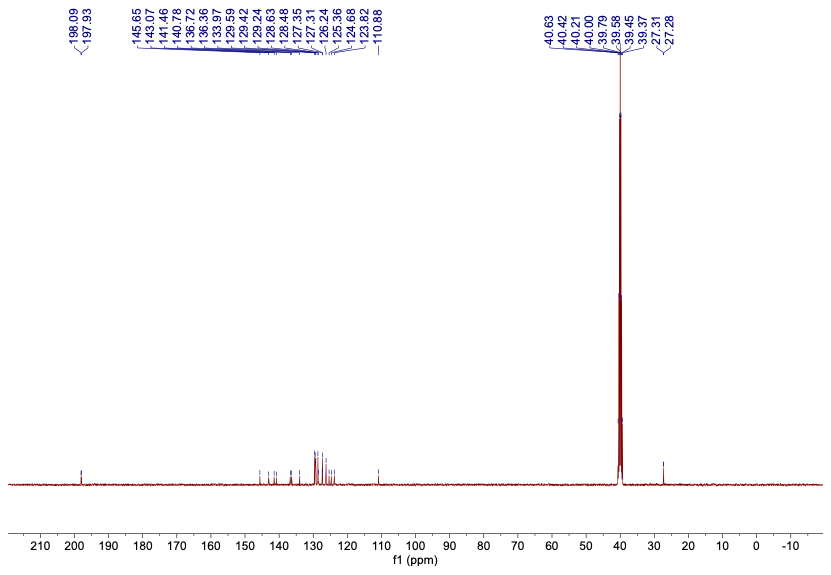

**Supplemental Figure S1: ^1^H and ^13^C-NMR spectra of CLM296:** 1,1'-((2-phenylimidazo[1,2-a]pyridine-6,8-diyl)bis(4,1-phenylene))bis(ethan-1-one) (**CLM296**)

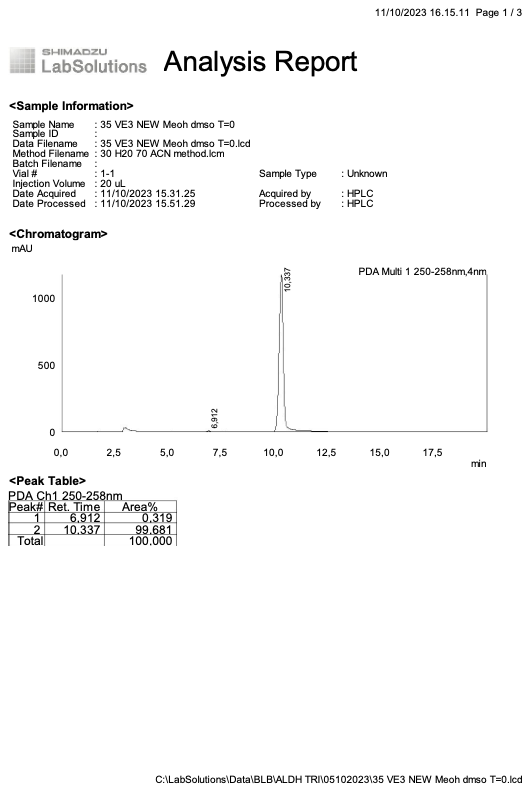

**Supplemental Figure S2:** **HPLC traces of CLM296.** *1,1'-((2-phenylimidazo[1,2-a]pyridine-6,8-diyl)bis(4,1-phenylene))bis(ethan-1-one) (****CLM296****)*

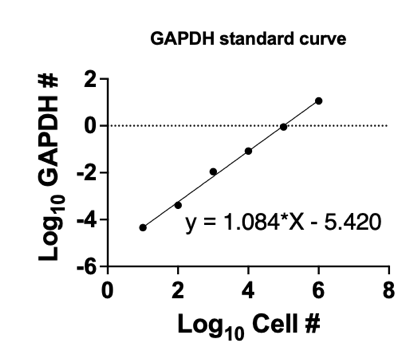

**Supplemental Figure S3. Human GAPDH standard curve spiked in naïve mouse lung lobes.**

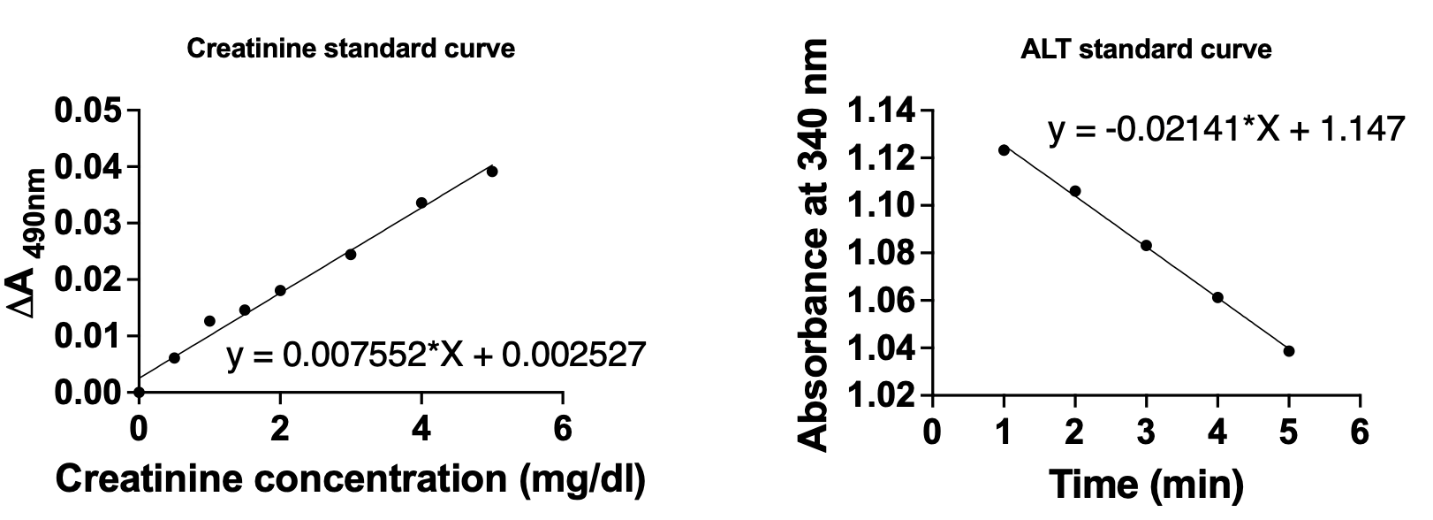

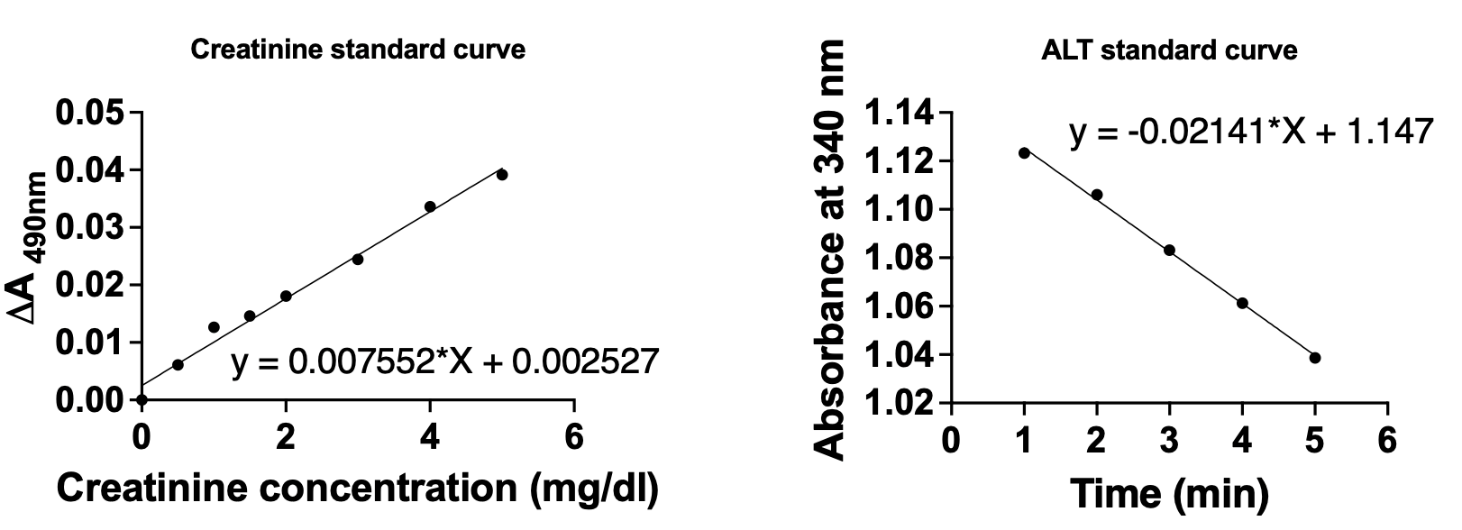
**A	B**

**Supplemental Figure S4. Standard curves for creatinine (A) or ALT (B) spiked in naïve mouse serum.**

**A**

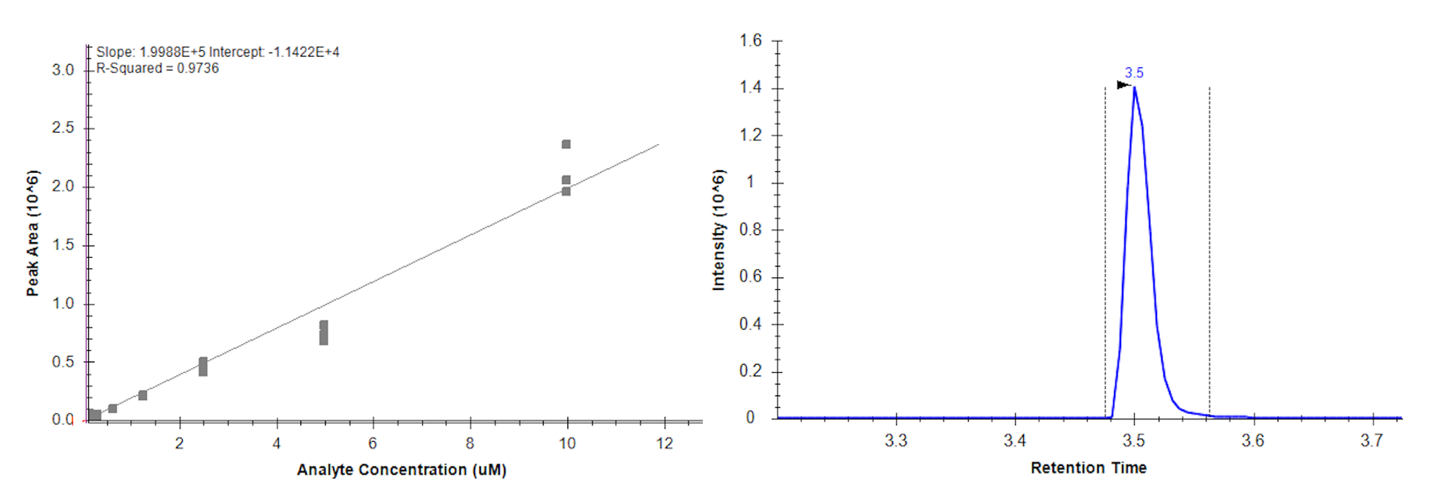

**B**

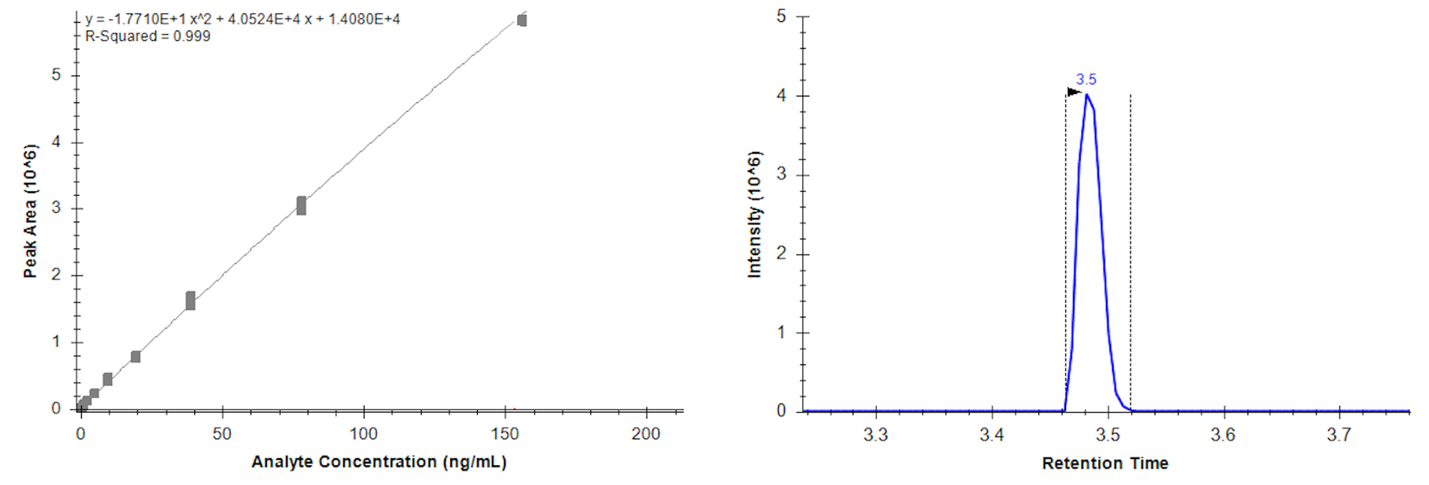

**Supplemental Figure S5. CLM296 and indomethacin quantification by LC-MS/MS.** **A**) Standard curve and retention time of CLM296 in 80:20 acetonitrile:acetone detected by LC-MS/MS. **B)** Standard curve and retention time of indomethacin in 80:20 acetonitrile:acetone detected by LC-MS/MS.

**A**

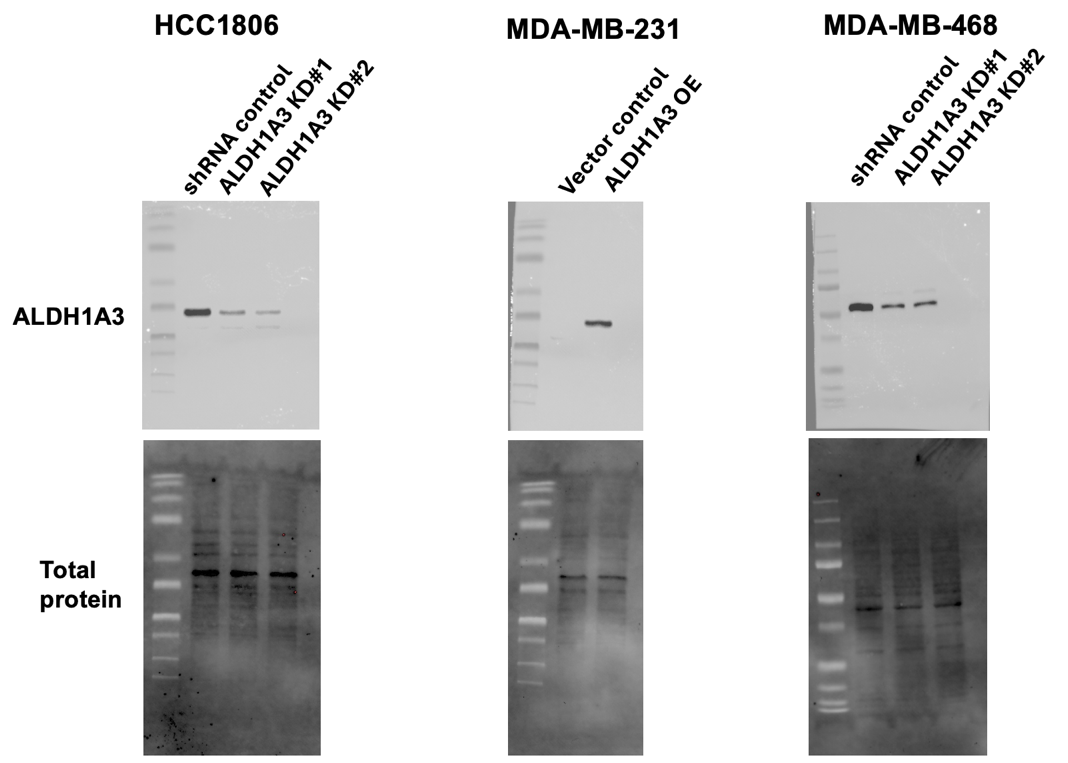

**B**

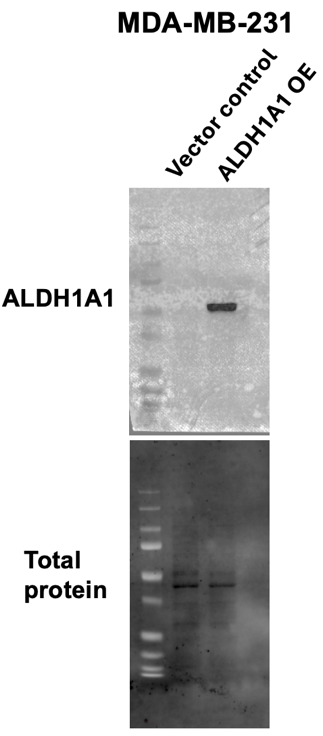

**Supplemental Figure S6. Uncropped Western blots. A)** ALDH1A3 western blots. **B)** ALDH1A1 western blot.

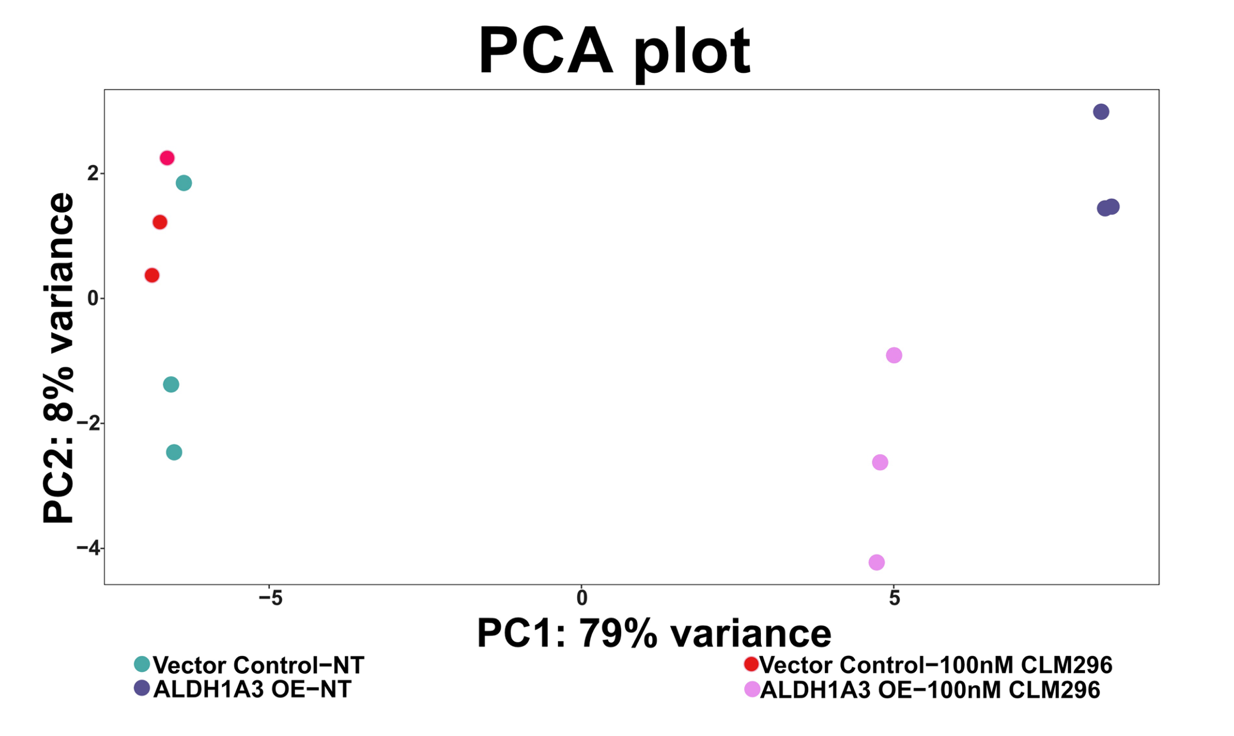

**Supplemental Figure S7. PCA plot showing variance in gene expression profiles of CLM296-treated or untreated vector control or ALDH1A3 OE MDA-MB-231 cells.**

**A**

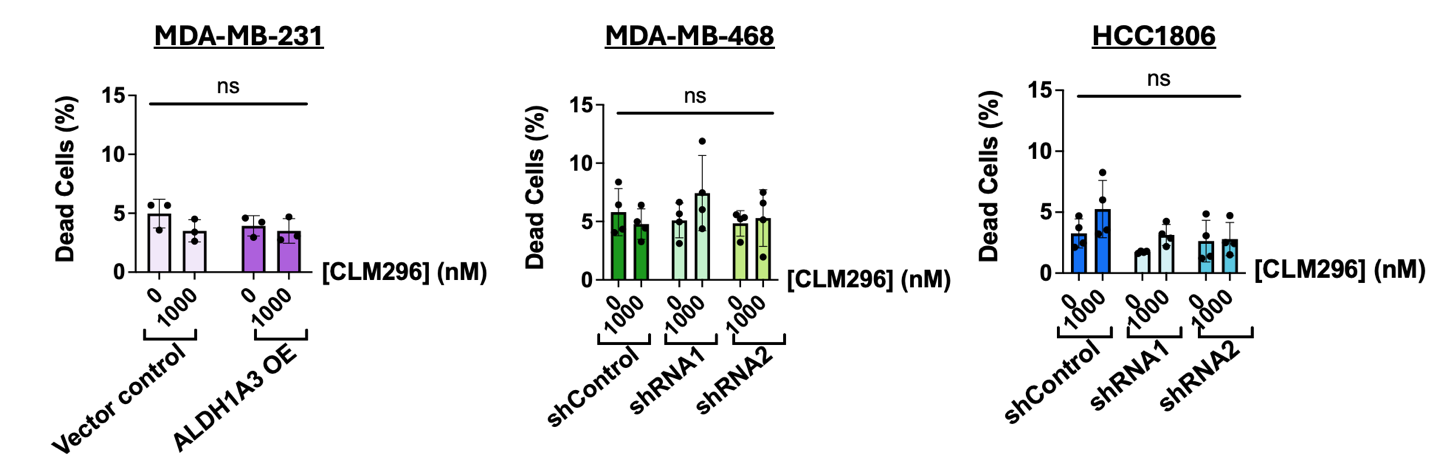

**B**
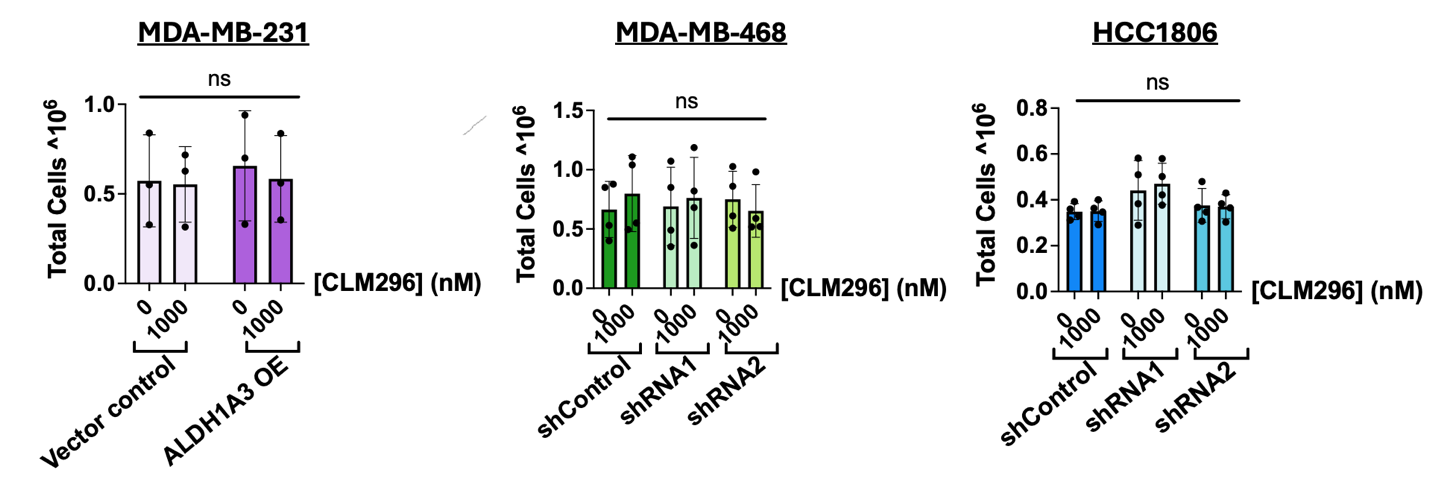

**Supplemental Figure S8. Neither ALDH1A3 nor 1µM CLM296 treatment for 72h affect cell viability and proliferation. A, B)** MDA-MB-231 (n = 3), MDA-MB-468 (n = 4) and HCC1806 (n = 4) cells with respective ALDH1A3 OE or ALDH1A3 KD clones treated with 1 μM CLM296 for 72 hours and total cells collected for quantification by cell counting with trypan blue staining. **A**) Percentage dead cells in each condition. **B)** Total cells. Each point represents a separate n, and error bars are represented as standard deviation. Significance determined by two-way ANOVA followed by multiple comparisons post-test and *p*-value < 0.05 = *, <0.01 = **, <0.001 = ***, <0.0001 = ****, ns = not significant.

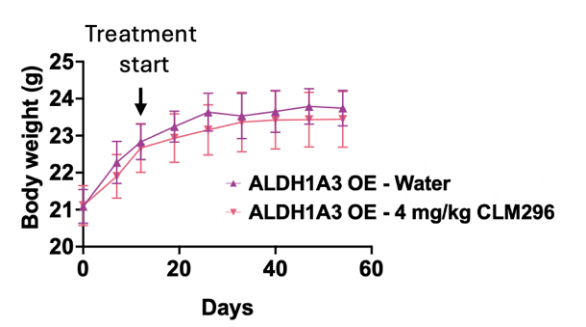
**A	B**

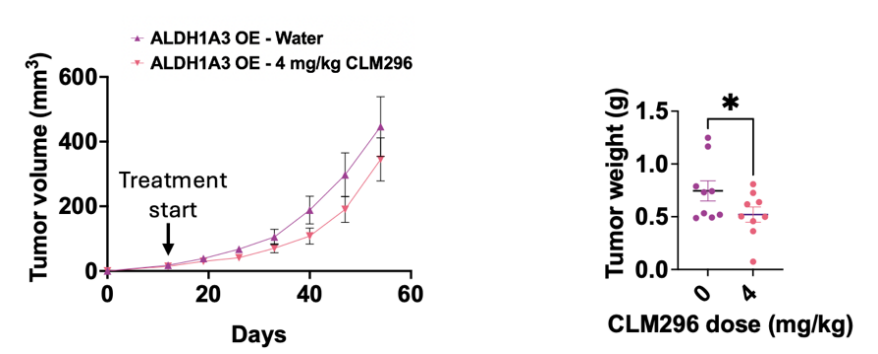

**Supplemental Figure S9. Tumor growth and mouse weights in the extended tumor study with MDA-MB-231 ALDH1A3 Overexpression, with or without CLM296 treatment. A)** Mouse weights (g), assessed weekly from the start of cell implantation. Error bars are represented as standard error mean. **B)** Tumor growth was assessed by tumor volumes and final tumor weights. Significance was calculated with unpaired, one-way *t-*test where *p*-value < 0.05 = *, and error bars are represented as standard error mean.

**SUPPLEMENTAL TABLES**

**Supplemental Table S1. qPCR primer sequences**

|  | **Gene** |  | **Sequence (5ʹ to 3ʹ)** |
| --- | --- | --- | --- |
| Reference Genes | **B2M** | Forward  Reverse | AGGCTATCCAGCGTACTCCA  CGGATGGATGAAACCCAGAC |
|  | **PUM1** | Forward  Reverse | GGCGTTAGCATGGTGGAGTA  CATCCCTTGGGCCAAATCCT |
|  | **RPL29** | Forward  Reverse | ACATGCGCTTTGCCAAGAAG  GGCTTAACCTCCTTGGGCTT |
|  | **ARF1** | Forward  Reverse | GTGTTCGCCAACAAGCAGG  CAGTTCCTGTGGCGTAGTGA |
|  | **TBP** | Forward  Reverse | GGCACCACTCCACTGTATCC  GCTGCGGTACAATCCCAGAA |
| Investigational genes | **ELF3** | Forward  Reverse | CCAGCGATGGTTTTCGTGAC GATGTCCCGGATGAACTCCC |
|  | **RAR**$\boldsymbol{\beta}$ | Forward  Reverse | GGTTTCACTGGCTTGACCAT GGCAAAGGTGAACACAAGGT |
|  | **DHRS3** | Forward  Reverse | TCTGTGATGTGGGCAACCG  ATGGTGATGTCACCCACCTTC |
|  | **Human GAPDH** | Forward  Reverse | TCAAGGCTGAGAACGGGAAG  CGCCCCACTTGATTTTGGAG |
|  | **Mouse GAPDH** | Forward  Reverse | GCGAGACCCCACTAACATCA  GGCGGAGATGATGACCCTTT |

**Supplemental Table S2. W/V ratio for pharmacokinetics study**

| **Organ** | **Weight/volume ratio** |
| --- | --- |
| Tumor | 5:3 |
| Liver | 5:2 |
| Lungs | 5:4 |
| Brain | 5:2 |
| Kidneys | 5:3 |
